# Culture of Saos-2 cells under hypoxic conditions stimulates rapid differentiation to an osteocyte-like stage

**DOI:** 10.1101/2023.04.23.537998

**Authors:** Anja R. Zelmer, Yolandi Starczak, Lucian B. Solomon, Katharina Richter, Dongqing Yang, Gerald J. Atkins

## Abstract

Few human osteocyte *in vitro* models exist and the differentiation of immature osteoblasts to an osteocyte stage typically takes at least 4-weeks of culture, making the study of this process challenging and time consuming. The osteosarcoma cell line Saos-2 has proved to be a useful model of human osteoblast differentiation through to a mature osteocyte-like stage. Culture under osteogenic conditions in a standard 5% CO_2_ and normoxic (21% O_2_) atmosphere results in reproducible mineralisation and acquisition of mature osteocyte markers over the expected 28-35 day culture period. In order to expedite experimental assays, we tested whether reducing available oxygen to mimic concentrations experienced by osteocytes *in vivo* would increase the rate of differentiation of Saos-2 cells. Cells cultured in a 5% CO_2_, 1% O_2_ atmosphere exhibited accelerated deposition of mineral, reaching near saturation by 14 days as demonstrated with the Alizarin Red and Von Kossa staining. The gene expression of the major hypoxia-induced transcription factor *HIF1α* and the key osteogenic transcription factor *RUNX2* were both elevated under 1% O_2_. Early (*COLA1, MEPE*) and mature (*PHEX, DMP1* and *SOST*) osteocyte markers were also upregulated earlier under hypoxic compared to normoxic growth conditions. Thus, culture under low oxygen accelerates key markers of osteocyte differentiation, resulting in a useful human osteocyte-like *in vitro* cell model within 14 days.

## 2. Introduction

Osteocytes are the major and most long-lived bone cell population, potentially living for decades *in vivo* (1). Osteocytes are critical for bone health and play numerous physiologic roles. The bone is a generally hypoxic tissue with oxygen levels varying between 1–6% O_2_ (2) thus osteocytes physiologically experience hypoxia *in vivo*. Hypoxia inducible factor-1 alpha (HIF-1α) is the principal transcription factor mediating adaptive responses to reduced O_2_ levels. HIF-1α protein levels are naturally regulated by proteosomal degradation following prolyl-hydroxylation by a family of prolyl-hydroxylases (PHDs1-3) (3). During hypoxia, prolyl-hydroxylation is blocked, leading to HIF-1α accumulation, nuclear translocation and dimerisation with HIF-1β, initiating HIF-responsive gene transcription by binding to hypoxia-responsive elements (HREs) in target gene promoters. Hypoxia signaling is an important regulator of normal bone mass, demonstrated by young *Hif1a*^*null*^ mice having reduced cortical bone volume, which is reversed in aged mice (4).

The human osteosarcoma cell line Saos-2 (or HTB-85™) has long been known to mineralise its extracellular matrix. We previously reported the ability of Saos-2 cells to differentiate in standard 2-dimensional (2D) cultures or in 3D cultures to an osteocyte-like stage by 28 days (28d) of osteogenic culture (5). The transition from an osteoblastic cell into an osteocyte-like cell can be monitored by the change in gene expression pattern and the increase in mineralisation. We recently showed the utility of these 28d cultures for the study of intra-osteoblastic and intra-osteocytic infection with *Staphylococcus aureus* (6). However, 28d of pre-culture prior to performing experimentation presents a logistical barrier and is costly in terms of both tissue culture reagents and time. We hypothesised that culture of these cells under more physiologic, low oxygen conditions, would promote their differentiation. In this study, we therefore compared osteogenic differentiation under atmospheric (normoxic) oxygen (21%) and a nominal hypoxic concentration of 1% O_2_.

## 3. Materials and Methods

### a. Cell culture

Saos-2 cells were maintained in growth media consisting of αMEM (Gibco, NY, USA) supplemented with 10% v/v foetal calf serum (FCS) and standard tissue culture additives (10 mM HEPES, 2 mM L-Glutamine, penicillin/streptomycin each 1 unit/ml (Thermo-Fisher, VIC, Australia)) at 37°C/5% CO_2_ (5-7). For experimentation, cells were seeded at a density of 1 × 10^4^ cells/well in 48 well plates or 2 × 10^4^ cells/well in 24 well plates and maintained with bi-weekly media change. To achieve an osteocyte-like phenotype, Saos-2 cells were switched to differentiation media at confluence, consisting of αMEM supplemented with 5% v/v FCS, standard tissue culture additives plus 50 µg/ml ascorbate 2-phosphate and 1.8 mM potassium di-hydrogen phosphate (Sigma, St Louis, USA) and then cultured either at 37°C/5% CO_2_ for 28 days in a standard atmosphere tissue culture incubator (Heracell Vios 160i, Thermo Fisher Scientific, Adelaide, SA, Australia) (5) or at 37°C/5% CO_2_/1% O_2_ in a nitrogen-controlled hypoxic incubator (New Brunswick Galaxy 170R, Eppendorf, Hamburg, Germany). Prior to media changes, medium for hypoxia cultures was equilibrated under 1% O_2_ to reduce dissolved oxygen levels. All experiments were performed in at least biological quadruplicates.

### b. Measurements of cell viability

Cells were cultured, as described above, on Cell Imaging Plates (Eppendorf, Hamburg, Germany). After 7-, 14-, 21-, and 28-days in differentiation media, the cells were incubated for 5 min with eBioscience™ Calcein Violet 450 AM Viability Dye (Invitrogen, Waltham, MA, USA) and Ethidium Homodimer III (Biotium, Fremont, CA, USA). Confocal images were taken with an Olympus FV3000 confocal microscope (Olympus, Tokyo, Japan) and processed with Fiji ImageJ to obtain the relative intensity.

### c. Measurement of in vitro mineralisation

Cells were harvested 7-, 14-, 21-, and 28-days after changing to differentiation media. They were rinsed with phosphate buffered saline (PBS), fixed with 10% formalin for 10 min, rinsed three times with milli-Q water (MQ). Four replicates were stained for calcium with 2% Alizarin Red for 5 min and rinsed with water until the wash was clear. Cells were imaged and then processed for quantitative analysis (5). For this, cells were incubated for 30 min with 10% acetic acid on a shaker and then transferred into 1.5 ml reaction tubes. After mixing the lysate was heated for 10 min at 85ºC and then put on ice for 5 min. The cold samples were centrifuged at 20,000g for 15 min and the supernatant transferred to fresh tubes. The pH was adjusted to 4.1-4.5 with 10% ammonium hydroxide. Samples were read at OD_405 nm_ and quantified using a standard curve. Four replicates were stained for deposited phosphate using the von Kossa stain (8). Briefly, cells were incubated with 1% AgNO_3_ for 30 min under light exposure, rinsed 3 times with MQ and incubated with Na_2_S_2_O_3_ for 5 min, rinsed 3 times with MQ and imaged.

### d. Measurement of alkaline phosphatase activity

Cells were harvested 7-, 14-, 21-, and 28-days after changing to differentiation media. They were rinsed with PBS, fixed with 10% buffered formalin for 10 min, then rinsed three times with MQ. Four replicates were stained for alkaline phosphatase (ALP) with StayRed (Abcam, Boston, MA, USA) according to the manufacturer’s instructions.

### e. Measurement of gene expression by real-time RT-PCR

For the quantification of gene expression of Collagen Type I (*COLA1*), Runt-related transcription factor 2 (*RUNX2*), bone gla-containing protein-1 (*BGLAP*), Hypoxia-inducible factor 1-alpha (*HIF1A*), matrix extracellular phosphoglycoprotein (*MEPE*), phosphate-regulating neutral endopeptidase (*PHEX*), dentin matrix protein 1 (*DMP1*) and sclerostin (*SOST*), total RNA was isolated using Trizol reagent (Life Technologies, NY, USA) and complementary DNA (cDNA) templates were prepared using the iScript RT kit (BioRad, CA, USA), as per manufacturer’s instructions. Real-time RT-PCR reactions were then performed to determine the mRNA levels of target genes using RT2 SYBR Green Fluor qPCR Mastermix (Qiagen, Limburg, Netherlands) on a CFX Connect Real Time PCR System (BioRad). The sequences of the oligonucleotide primer sets targeting each gene are listed in **Table 1**. Gene expression relative to the level of *ACTB* mRNA was calculated using the 2^−ΔCt^ method.

**Table 1:**
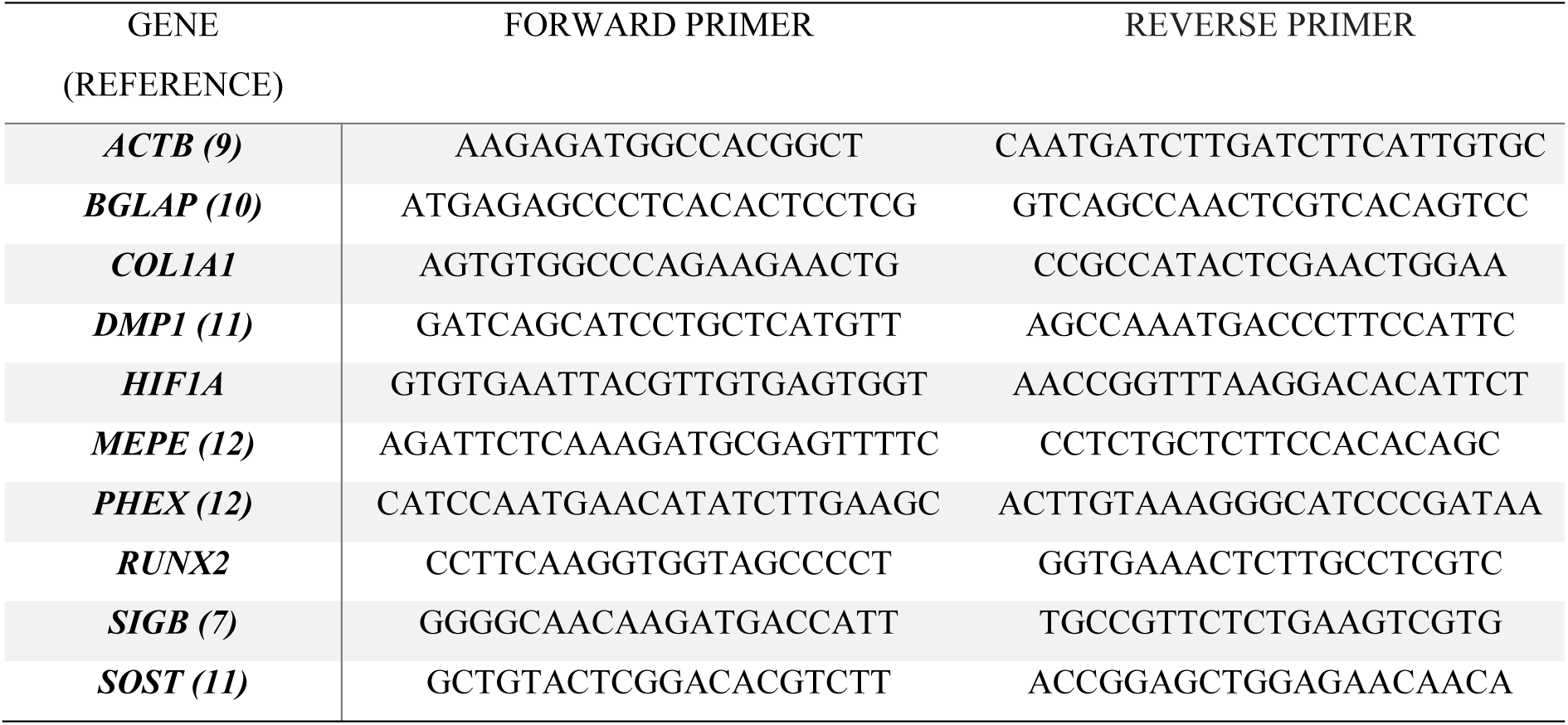
Oligonucleotide primer sequences used for qPCR

### f. Statistical analysis

Two-way ANOVA with Tukey’s post-hoc tests were used to compare differences during the time course of Saos-2 differentiation. To compare specific pairs, T-tests were used. All analysis was performed using GraphPad Prism 9 software (GraphPad, CA, USA). Values for p < 0.05 were considered significant.

## 4. Results

### a. Effects of oxygen level on cell viability

At all time-points examined, cells under either normoxic or 1% O_2_ culture condition appeared uniformly viable by phase contrast microscopy and by Calcein-Violet and Ethidium Homodimer III staining (**Fig. 1A**). Live/dead staining confirmed that the proportion of dead cells did not significantly change during the differentiation, nor was it different under either culture condition, with at least 85% viability throughout (**Fig. 1B**), indicating that Saos-2 cells can survive under 1% O_2_ levels as well as under normoxia for at least 28 days.

**Figure 1:**
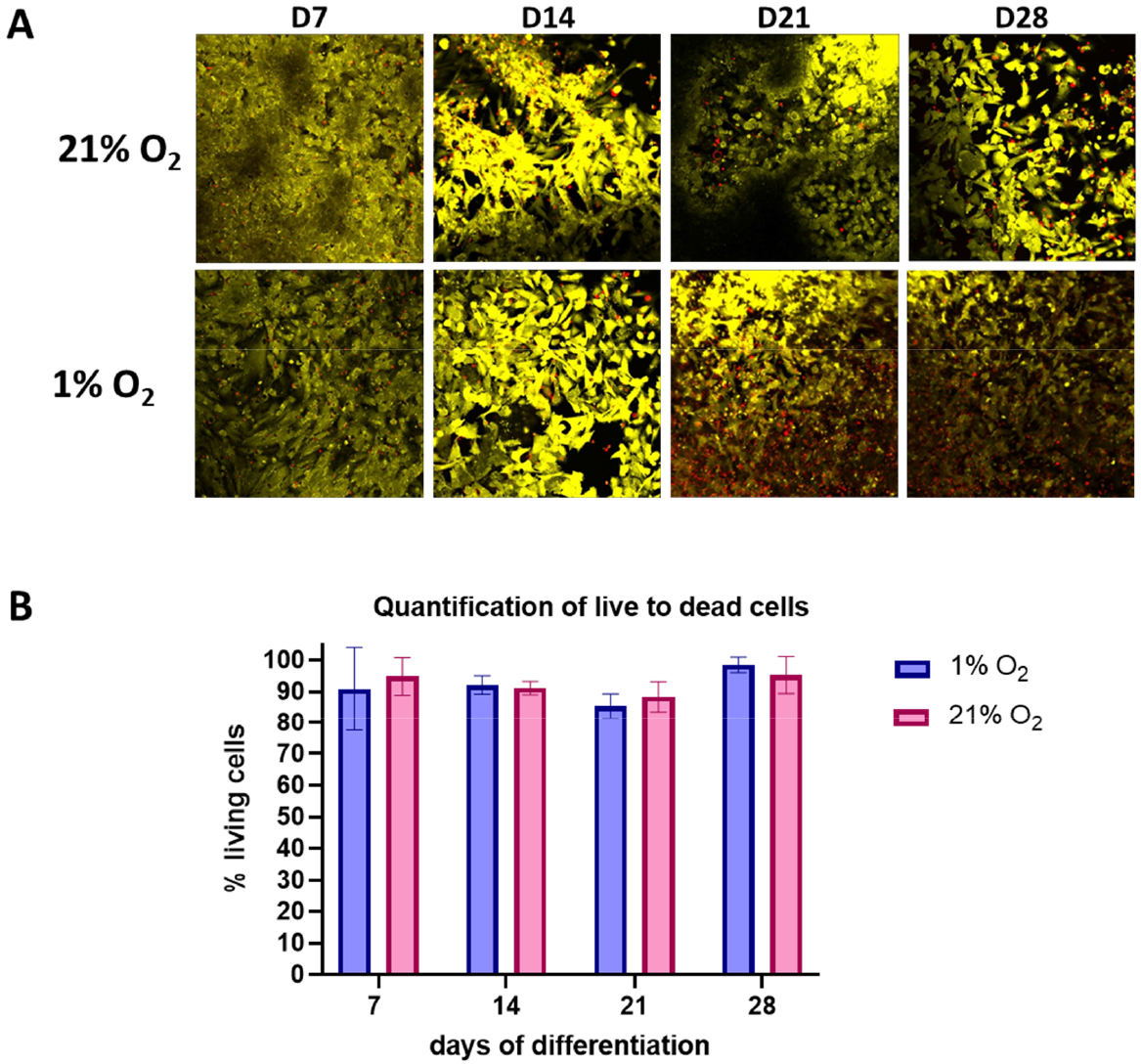
Saos-2 cell after 7-, 14-, 21-, and 28-days of differentiation under 1% (blue) or 21% O2 (pink) stained with Calcein Violet 450 AM Viability Dye and Ethidium Homodimer III for live/dead confocal imaging. A) Representative sample image B) quantified percentage of live to dead stains.

### b. Effects on in vitro mineralisation

Saos-2 cultures differentiated under 1% O_2_ mineralised significantly more rapidly than under 21% O_2_, as determined by Alizarin Red staining for calcium and von Kossa staining for phosphate (**Fig. 2**). Cells under 1% O_2_ reached a significantly higher mineralisation level at 14 days than the cells under 21% O_2_ did after 28 days and was maximal at the 14d time point (**Fig. 2B**). Consistent with this, under 1% O_2_ more active ALP was detectable than under 21% O_2_ (**Fig. 2C**).

**Figure 2:**
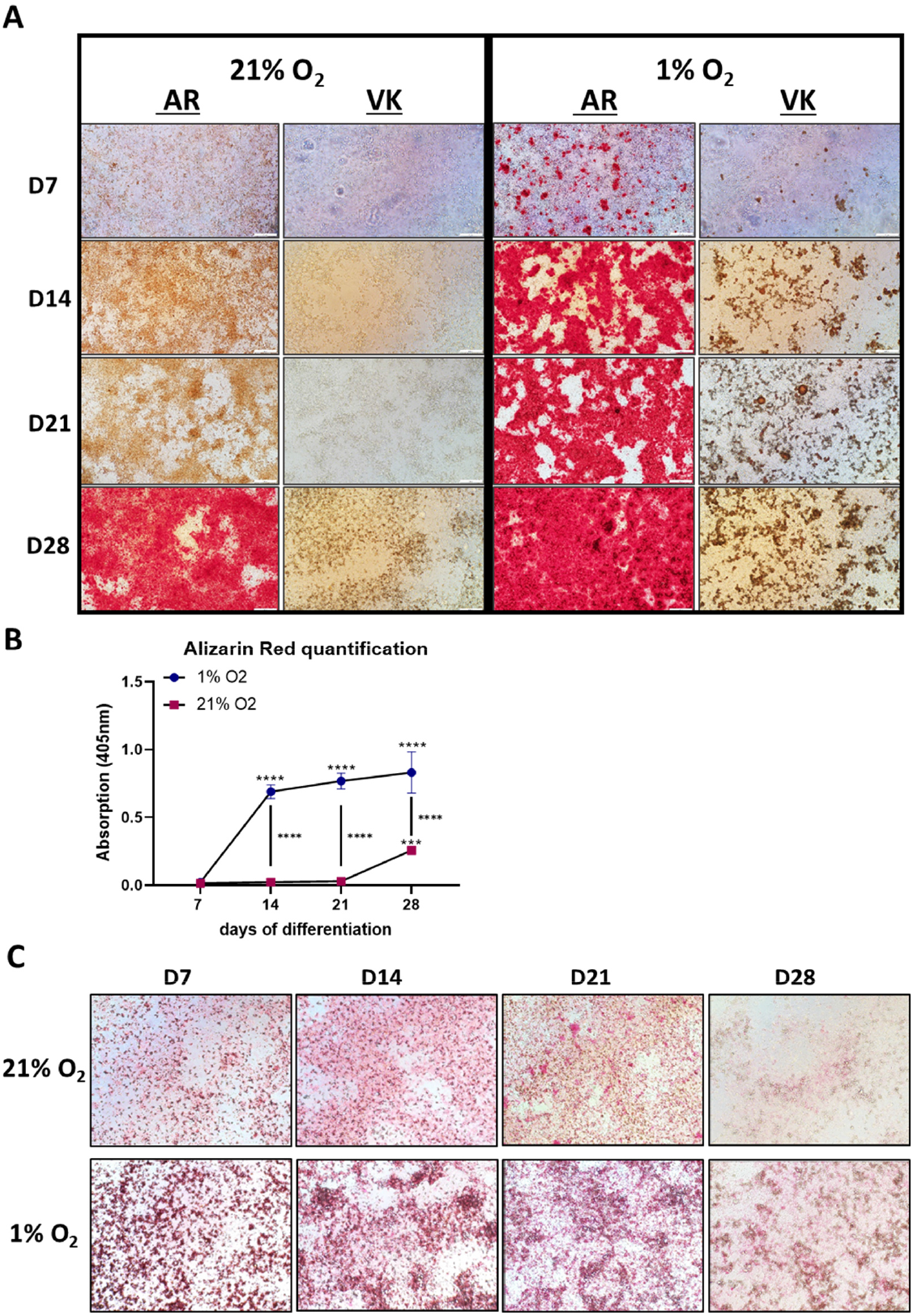
Saos-2 cell after 7-, 14-, 21-, and 28-days of differentiation under 1% or 21% O2 A) stained with Alizarin Red (AR) for calcium and von Kossa (VK) for phosphate to cell induced observe mineralisation. B) Alizarin Red quantification at 405 nm absorption. Asterisk above a time-point indicate significant difference to d7. C) stained with StayRed (Abcam) to measure alkaline phosphatase activity.

### c. Effects on gene expression

There was a significant higher expression of *HIF1A* mRNA under 1% O_2_ compared to 21% O_2_, however only from day 14 of culture (**Fig. 3A**), consistent with the cells responding metabolically to the low oxygen conditions. Expression of the major osteoblastic transcription factor *RUNX2* was elevated from day 14 under 1% O_2_, (**Fig. 3B**) consistent with hypoxia having an osteogenic effect on the cells (13). The expression of *COL1A1* increased under both culture conditions, as expected but to a greater extent under 1% O_2_ (**Fig. 3C**), indicating increased type I collagen bone matrix production, consistent with the increased mineralisation under hypoxic conditions. The expression of late osteoblastic/osteocytic markers *BGLAP, MEPE* and *PHEX* (**Fig. 3D-F**) followed a similar pattern under both culture conditions, and at 21d enhanced under 1% O_2_ in the case of *MEPE* and *PHEX*, consistent with at least retention, or enhancement of osteogenic differentiation under a hypoxic stimulus. Elevated mRNA expression of the mature osteocyte markers *DMP1* and *SOST* throughout differentiation was expected (5, 14-17) but these were both relatively increased under 1% O_2_ by day 14 (**Fig. 3G-H**), indicating more rapid acquisition of an osteocyte-like phenotype than under normoxic conditions.

**Figure 3:**
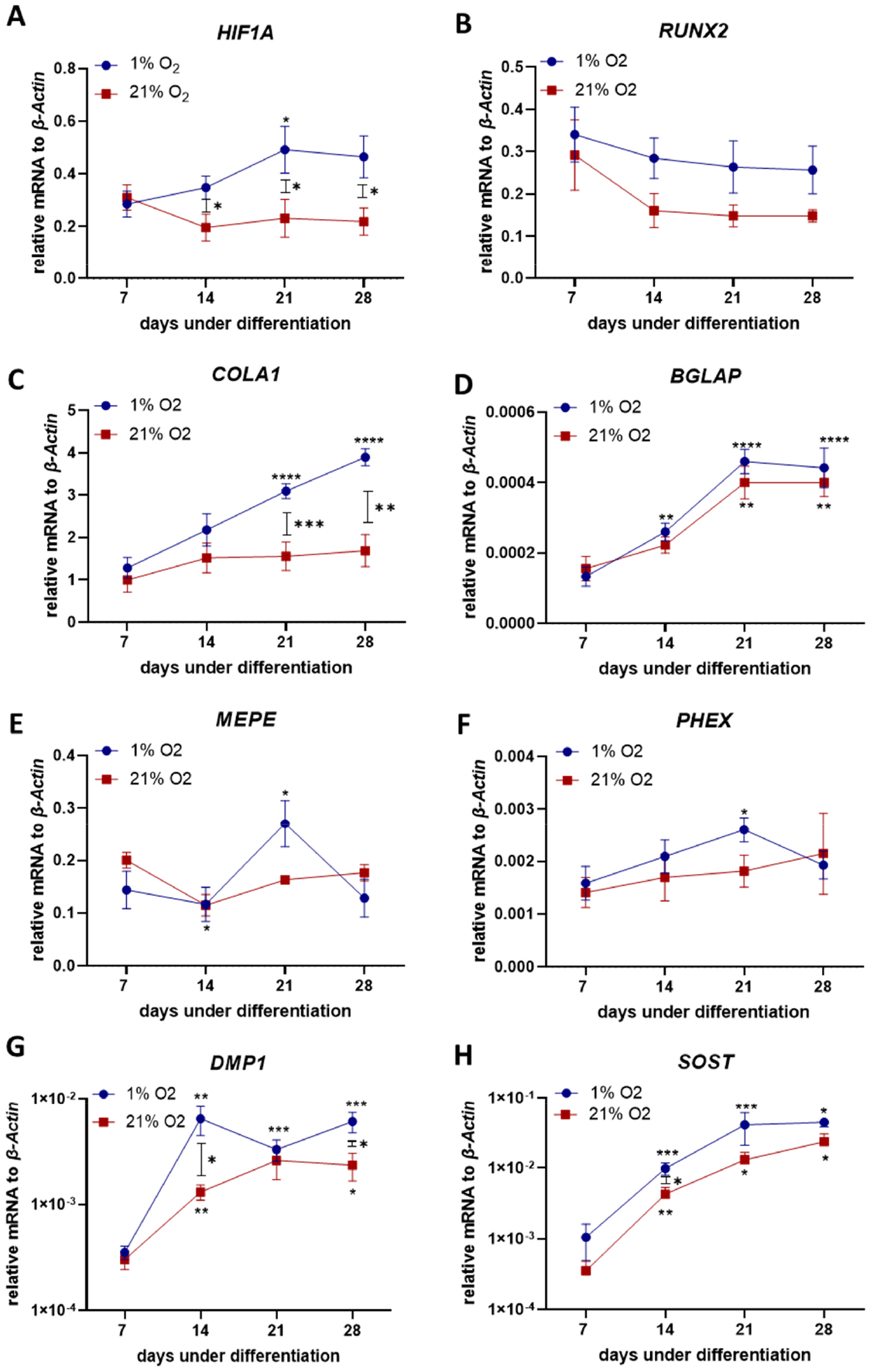
mRNA expression of osteocyte markers in Saos-2 cell after 7-, 14-, 21-, and 28-days of differentiation under 1% (blue circle) or 21% O2 (pink square): A) *Hif1A*, B) *RUNX2*, C) *Col1A1*, D) *BGLAP*, E) *MEPE*, F) *PHEX*, G) *DMP1* and H) *SOST*.

When comparing the 14d hypoxic time-point to the 28d normoxic time point, *PHEX, DMP1* and *SOST* were at least at the same expression level, indicating that under hypoxic conditions, cells express similar levels of osteocyte marker genes after 14 days as cells under normoxic conditions do after 28 days of differentiation.

## 5. Discussion

Saos-2 cells are the only widely validated transformed human osteoblastic cell line demonstrated to differentiate to an osteocyte-like stage (5). To date, one major limitation of such differentiation models is the long period of time, typically 28-35 days, required before the cells are ready to test as an osteocyte-like cell; here, we showed that differentiation under 1% oxygen reduces the differentiation period to 14 days, permitting more timely execution of these experiments.

Culture under 1% O_2_ induced higher expression of *HIF1A* mRNA compared to 21% O_2_ from day 14 of culture, consistent with the cells responding metabolically to the reduced available oxygen. Interestingly, there was no change in *HIF1A* levels at the day 7 time point under 1% O_2_. It is possible that because oxygen levels were not controlled during media changes beyond prior equilibration at 1% O_2_ that the hypoxic effect was slower to manifest or that the early undifferentiated cells were naturally adapted to a low oxygen environment; in either case, it is also possible that the extensive mineralisation by day 14 further reduced oxygen availability to the cells and this was an additional trigger for *HIF1A* upregulation. Regardless, the accelerated differentiation process under 1% O_2_ was reflected in more rapid *in vitro* mineralisation, indicated by maximal Alizarin Red staining by day 14 as a measure of calcium incorporation, increased ALP activity, permissive for phosphate incorporation, and elevated *COL1A1* mRNA expression as a measure of earlier collagen type 1 organic matrix production.

*RUNX2* expression by immature osteoblasts is fundamentally permissive for their osteogenic differentiation (13, 18), thus the elevated expression of *RUNX2* mRNA under 1% O_2_ compared to standard normoxic conditions is strongly supportive of an osteogenic response. This was reflected in the associated increase in expression of all osteogenic genes examined. *PHEX* and *MEPE* are classic markers for the osteoid osteocyte, or mineralising osteocyte, stage (19, 20), which slightly increased over time under normoxic conditions but peaked after 21 days under hypoxic conditions, consistent with an increased rate of differentiation. In similar previous experiments (under normoxia), both genes were shown to peak relatively late in primary human osteoblast and Saos-2 cultures during differentiation (5, 21). Most importantly, the mature osteocyte markers *DMP1* and *SOST* (14, 15, 22) increased significantly under both conditions but were relatively significantly increased under hypoxic conditions by day 14, and at this time point reached a similar expression level as after 28d under normoxic conditions. This is consistent with Saos-2 cells under hypoxic conditions reaching an osteocyte-like stage within 14 days.

Hypoxia has been shown to have profound effects on bone cell metabolism and osteoblast differentiation, although with sometimes contradictory findings (23). In primary rat calvarial osteoblasts, a hypoxia-dependent decrease in mineralised nodule formation was found, associated with decreased alkaline phosphatase activity (24), which is clearly the opposite to the effects observed in our study. Another study showed reduced *SOST* expression and increased Wnt/β-catenin signalling in the rat UMR106.01 osteosarcoma and mouse MLO-A5 osteoblastic cell lines, which like Saos-2 are both long bone-derived, consistent with a pro-anabolic effect of hypoxia, although effects on *in vitro* mineralisation were not reported (25). However, in our study *SOST* expression was further increased under 1% O_2_, we suggest as a physiological response to the increased mineralisation and acquisition of a mature osteocyte-like phenotype. The reasons for the observed differences between these studies and our own are unclear. It is possible that the effect of hypoxia may either be cell type (*e*.*g*. species, skeletal origin, transformed *v*. primary) or methodology-specific, as the above studies also employed different approaches to model hypoxia.

This modified Saos-2 differentiation model has a number of applications in terms of improved feasibility and physiologic relevance. Notably, studies investigating human osteoblast (Saos-2) on-growth onto modified orthopaedic implant surfaces, including those from our own group (26, 27), have typically utilised standard normoxic conditions, however the interface between an implant and the bone is usually avascular and therefore hypoxic. Another clear application is in terms of studying infection by pathogens such as *Staphylococcus aureus*. We have shown that osteocyte-like Saos-2 cells can internalise and harbour *S. aureus* (6). Further, since it has been observed that *S. aureus* adapts to hypoxic conditions (28, 29), it may be paramount to use hypoxic conditions for *in vitro* assays of *S. aureus* in osteomyelitis. In addition, Saos-2 cells are a useful cell line model, with which to study the gene and protein expression regulation of *SOST*/sclerostin in a human context, for example in response to BMP stimulation,(30) mechanical loading (31) and 1,25-dihydroxyvitamin D_3_ (32, 33), and it is possible that examination under hypoxic conditions would increase the physiological relevance of such studies.

In summary, Saos-2 cells cultured under low oxygen exhibit accelerated *in vitro* mineralisation and differentiation to a mature osteocyte-like stage, and achieve a useful osteocyte-like phenotype within 14 days. This model significantly reduces the time and therefore costs required to generate mature human osteocyte-like cultures, facilitating research into this important cell type. The model continues the advantage of using Saos-2 cells due to their ease of availability over primary cells, and improved intra-assay consistency independent of human donor variation. As a human cell line, the model may also provide findings of closer immediate clinical relevance than the extant non-human *in vitro* models.

## 6. Acknowledgements

ARZ was supported by University of Adelaide Faculty of Health and Medical Sciences Postgraduate Research Scholarships. This work was supported by funding from the National Health and Medical Research Council of Australia (NHMRC; Grant No. 2011042).

